# Crystal structure of arginine-bound lysosomal transporter SLC38A9 in the cytosol-open state

**DOI:** 10.1101/288233

**Authors:** Hsiang-Ting Lei, Jinming Ma, Silvia Sanchez Martinez, Tamir Gonen

**Affiliations:** Janelia Research Campus, Howard Hughes Medical Institute, 19700 Helix Drive Ashburn VA USA; Howard Hughes Medical Institute, University of California, Los Angeles, Los Angeles CA 90095 USA; Departments of Physiology and Biological Chemistry, David Geffen School of Medicine, University of California, Los Angeles, Los Angeles CA 90095 USA

## Abstract

Amino acid-dependent activation of mechanistic target of rapamycin complex 1 (mTORC1) is essential to reflect nutrient availabilities in cells for cell growth and metabolism^1^. Solute carrier 38 family A member 9 (SLC38A9) is the lysosomal transporter responsible for amino acid sensing in the mTORC1 signaling pathway^2–4^. Here we present the first crystal structure of SLC38A9 from *Danio rerio* in complex with arginine. As captured in the cytosol-open state, the bound arginine was locked in a transitional state stabilized by the transmembrane helix 1 (TM1) of SLC38A9 which was anchored at the grove between transmembrane helix 5 and 7 inside the transporter. The key motif WNTMM on TM1, contributing to the anchoring interactions, is highly conserved in various species. Mutations in WNTMM motif abolished arginine transport by SLC38A9. The underlying mechanism of substrate binding is critical for both sensitizing mTORC1 signaling pathway to amino acids and for maintaining amino acid homeostasis across lysosomal membranes^2^.

---

SLC38A9 is a sodium-coupled transporter that transports a variety amino acids across lysosomal membranes ^3,5^. Classified within Amino Acid-Polyamine Organocation (APC) superfamily^6^, SLC38A9 belongs to the Amino Acid/Auxin Permease (AAAP) subfamily^7^. To date, members of AAAP family are found only in eukaryotic systems and share features like extended N-terminal soluble domain and 11 transmembrane helices^7,8^. In *Homo sapiens*, at least 17 known AAAP members were found ^9^, spanning across solute carrier families SLC32, SLC36 and SLC38, although none of these structures have been determined to date. Thus far, 11 members of SLC38 have been identified in humans^10^, two of which, SLC38A7 and SLC38A9, localize at lysosomes ^5,11,12^. SLC38A9 is the only member of this family known to participate in the Ragulator-Rag GTPases complex and in turn plays an important role in the amino acid dependent activation of mTORC1 ^3,5,13–15^.

It is unclear how the SLC38A9 senses amino acids and how it transmits the signals to affect the mTORC1 pathway, although previous studies showed that two parts of SLC38A9, its N-terminal domain and its transmembrane domain, are responsible for distinct functions and make up a complete amino acid sensor to mTORC1 signaling pathway^3,5^. The N-terminal domain of SLC38A9 directly interacts with Rag GTPases and Ragulator. It has been reported that over-expression of this N-terminal domain in HEK-293T cells resulted in an activation of mTORC1^3,5^. On the other hand, the transmembrane domain of SLC38A9 functions as an amino acid sensor and when in amino acid deficient environment it lowers the activity of mTORC1 ^2,3,5^. Like many secondary transporters, the transmembrane domain of SLC38A9 likely switches between various conformational states during transport^16–18^ and it is possible that these conformational changes affect the structure of the N-terminal domain and hence the interactions between SLC38A9 and Ragulator or Rag GTPases ^2,3,5^.

In the present study, we determined the crystal structure of SLC38A9 from zebrafish (*Danio rerio*, drSLC38A9) in complex with arginine at 3.17 Å (Fig. 1a, 1b, Table S1). The transporter consists of 11 transmembrane (TM) helices with its N-terminus located in the cytosol while its C-terminus on the lumen side (Fig 1a). Consistent with other members in the APC super family^9^, SLC38A9 adopts a LeuT-like pseudo-symmetric bundle of five transmembrane-helix inverted-topology repeat: the N-terminal half consists of TMs 1-5 and the C-terminal half consists of TMs 6-10. Transmembrane helix 11 flanks the transporter from one side. TMs 1 and 6 are broken and line the substrate binding where an arginine molecule was identified (Fig. 1b). Sequence alignment of this protein from zebrafish, frog, mouse and humans indicate that the eleven transmembrane regions are highly conserved (Fig. S1). drSLC38A9 shares 61.9% of identity and 86.6% of similarity with the human homolog (hSLC38A9) (Fig. S1). Despite significant efforts to crystallize full-length drSLC38A9, only the N-terminal truncated form (ΔN-drSLC38A9) yielded ordered crystals amenable to diffraction studies. An antibody fragment, Fab-11D3, was used to further stabilize the luminal loops and further optimize the crystallization (Fig. S2, S3). As shown by the arginine uptake assay using SLC38A9-reconstituted liposomes (Fig. 1c, S4), the truncated SLC38A9 is active and able to bind and transport arginine as the wildtype protein.

**Figure 1.**
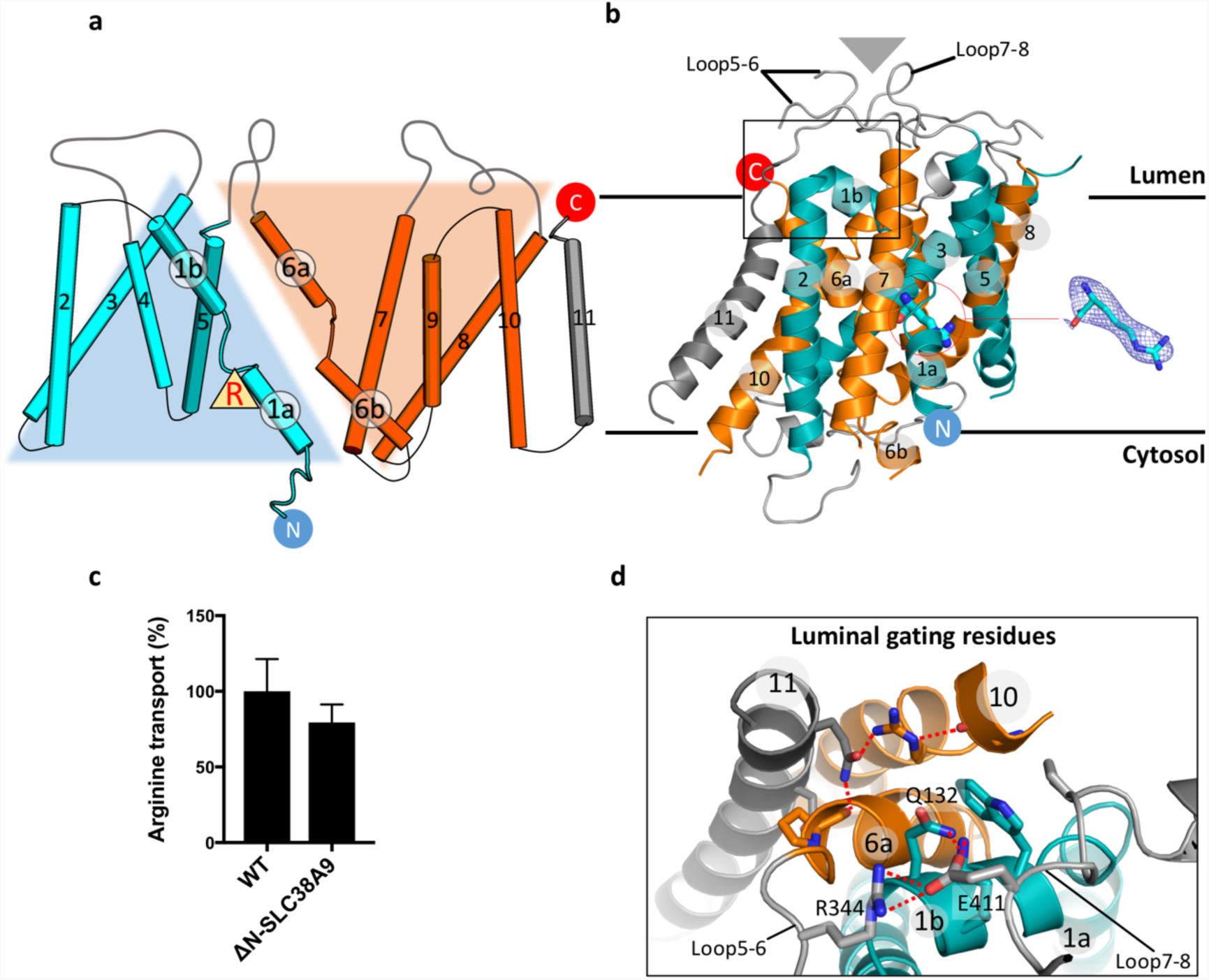
Overall architecture and the luminal gating network of arginine-bound SLC38A9. **a**, Two-dimensional topology model of SLC38A9. The first ten transmembrane helices are folded into a characteristic 2-fold LeuT-like pseudo-symmetry (five transmembrane-helix inverted-topology repeat). Bound Arg is marked by a filled yellow triangle, next to the TM1a helix. Cyan, TM1-5; orange, TM6-10; grey, TM11. **b**, SLC38A9 structure on lysosomal membranes. Transmembrane helices are colored as in **a**. Position of Fab fragment is shown by grey triangle above the luminal loops. Lumen and cytosol domains are equivalent to extracellular (Out) and intracellular (In) domains in a transporter expressed on cell plasma membranes. An Arg (blue stick) is identified at the binding site next to TM1a. 2mFo-DFc map contoured at 1.0 σ is shown for the Arg. **c**, SLC38A9-reconstituted liposome-based uptake assay shows that the efficiency of arginine transport is similar in wildtype and the truncated SLC38A9 used for crystallization. Percent transport is calculated by normalizing the mean of WT to 100 %. Error bars represent standard error of the mean (s.e.m.) of triplicate experiments. **d**, Enlarged view of the boxed region in **b** encompassing the luminal gating residues. Glu 411 of Loop7-8 act as a network hub in the luminal gating of SLC38A9, joined by Lys 131, Gln 132 of TM1b and Arg 344 of Loop5-6.

Because SLC38A9 is found in the lysosomal membrane, its cytosol-open state resembles an inward-open state for a transporter found in the cell plasma membranes (Fig. 1b). In the cytosol-open state presented here, the luminal gate is closed while the cytosol side consists of a wide vestibule open to the cytoplasm (Fig. 1d and Fig. 2). At the luminal side, polar interactions between residues on TM1b, Loop5-6, Loop7-8 and residues on TM6a, TM10, TM11 prevent solvent access to the substrate binding site toward the center of the transporter (Fig. 1b and d). Unlike other APC transporters (such as LeuT, BetP, CaiT, MhsT) in their inward-open state^19–21,22^, SLC38A9 does not use an inner gating system at the central region to keep solvent out. In contrast, it closes its luminal surface by the peripheral polar groups, Lys 131, Gln 132 on TM1b, Arg 344 on Loop 5-6 and Glu 411 on Loop7-8 (Fig. 1d).

**Figure 2.**
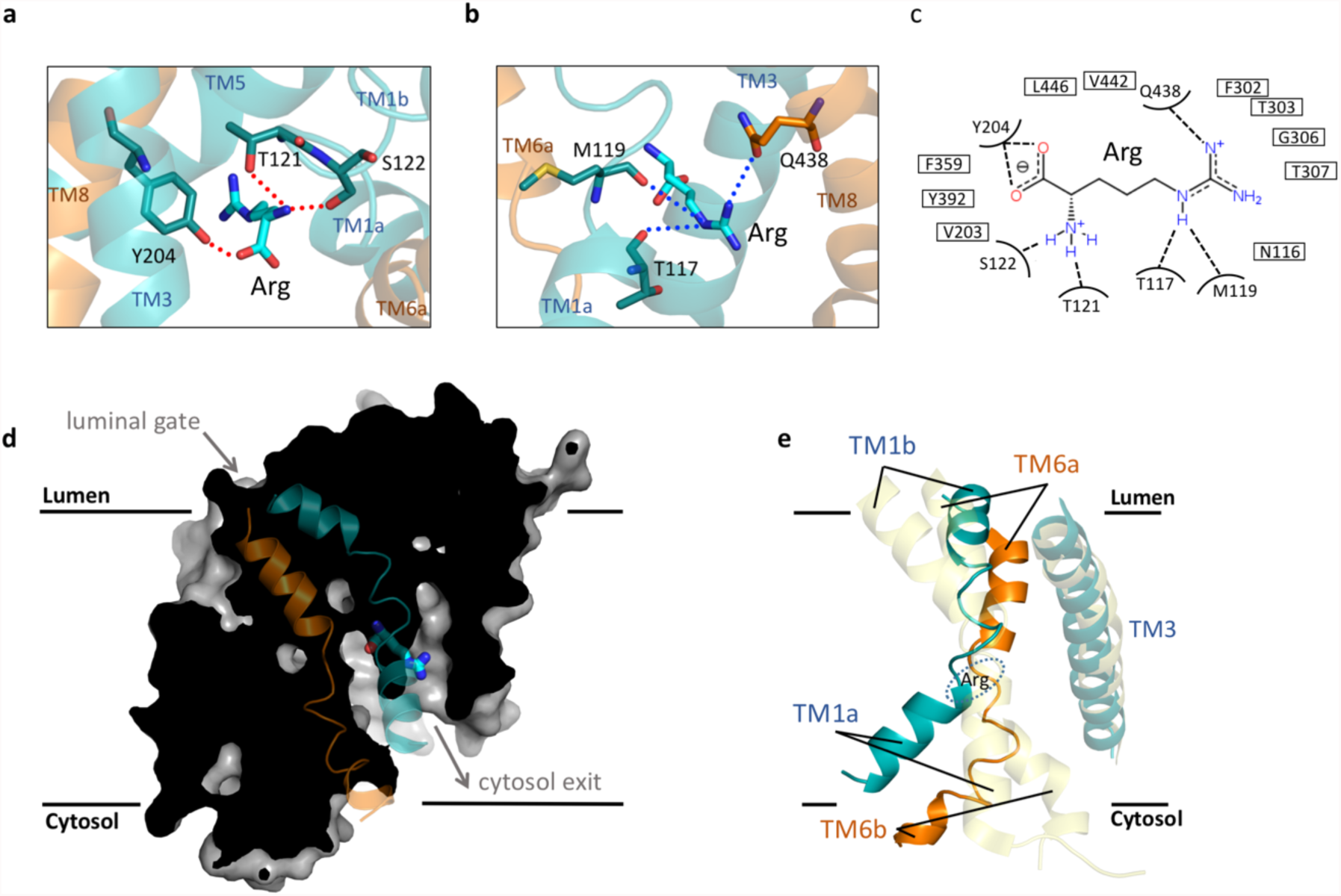
Arginine bound in the cytosol-open conformation of SLC38A9. **a**, View from the carboxylate group of Arg. The TM1a binding site is surrounded by TM1a, TM3, TM6b, TM5 and TM8. α-amino group of Arg is bonded to β-hydroxyl group of Thr 121 and Ser 122. Carboxylate group of Arg is bonded to Tyr 204. **b**, View from the guanidinium group of Arg. The positive group is stabilized by Thr 117, Met 119 on TM1a and Gln 438 on TM8. **c**, Schematic representation of the SLC38A9 and Arg interactions shown in panel **a** and **b**. Dotted lines depict the hydrogen bonds. **d**, Surface presentation of SLC38A9 in the cytosol-open state. TM1 and TM6 are colored in cyan and orange, respectively. **e**, Superposition of SLC38A9 (colored as in Fig. 1a) and AdiC (3L1L)^23^ in translucent yellow. TM3, 4, 8, 9 are used in the superimposition. Location of the bound Arg in SLC38A9 is shown by dashed oval. Structural alignment is performed in PyMOL. AdiC to SLC38A9; RMSD=3.57 Å, Cealign for 72 residues.

SLC38A9 was crystallized in the presence of its substrate arginine. The electron density map allowed us to identify an arginine molecule bound close to the center of SLC38A9 adjacent to transmembrane helix 1 a (TM1a) (Fig 1b and Fig. 2a-d). Recognition of arginine at this location involves interactions with Thr 117, Met 119, Thr 121, Ser 122 from TM1a, Tyr 204 on TM3 and Gln 438 from TM8. The α-amino group of arginine is hydrogen bonded to Thr 121 and Ser 122, and is further stabilized by the 4-hydroxyl group of Tyr 204 across the cytoplasmic vestibule on TM3 (Fig. 2a). Notably, a surface area made from the backbone carbonyl groups of Asn116, Thr 117, Met 118 and Met 119 electrostatically draw the guanidinium group of the bound arginine adjacent to TM1a (Fig. 2b). It has been shown that in the human homolog, the mutation T133W (equivalent to T121W in the present structure) abrogates transport activity for arginine^2^. Indeed, in our structure Thr 121 is a key residue involved in stabilizing the bound arginine.

Superposition of SLC38A9 with other related transporters that were captured at different states may provide insights for the mechanism of transport and for the structural reconfigurations that result from the transition from one state to another. One useful comparison is the arginine/agmatine antiporter AdiC^23^ which shares the same fold as SLC38A9 but was captured in a different state. Although AdiC does not couple sodium ions when transporting substrates, the location of the central site in AdiC resembles that of LeuT, a model system for a Na+-coupled transporter (Fig S5). In AdiC, the α-carboxyl and -amino group of arginine are bound between TM1 and TM6 while the guanidinium side group of the arginine was found next to TM3. Conventionally, TM3, 4, 8, 9 are defined as the scaffold domain in LeuT-like APC transporters because minimum changes were found in structures at different states ^17^. By superposing the scaffold domain of AdiC to SLC38A9, the change in the relative positions of TM1 and TM6 between the two transporters provide clues of the possible transformations between lumen-open state and cytosol-open state (Fig. 2e). The positions of TMs 1 and 6 change significantly during the transport cycle and their movement resembles a ratchet-like movement like what was observed in other LeuT-like APC transporters^24^.

As TMs 1 and 6 change conformation they affect the location and bonding of arginine. The arginine in SLC38A9 was found at a different location than the arginine from AdiC (Fig 3a). In AdiC, an arginine molecule was observed at the center of the transporter lying roughly parallel to the plane of the membrane. However, in SLC38A9 which is open to the cytosol, the arginine changed its orientation to point toward the cytosol (Fig 3a). This orientation of the arginine is stabilized by interactions with TM1a. The binding at TM1a site requires specific geometry of the amino acid, such as an elongated, positively charged side group. The location of the bound arginine in SLC38A9 is distinct from other substrates found in transporters that were also captured in the inward-open/occluded conformation, for instance, vSGLT, BetP, CaiT and MhsT^20–22,25^. For SLC38A9, the unique binding site of arginine may suggest that the present structure represents a divergent intermediate state in the transport cycle upon arginine binding.

**Figure 3.**
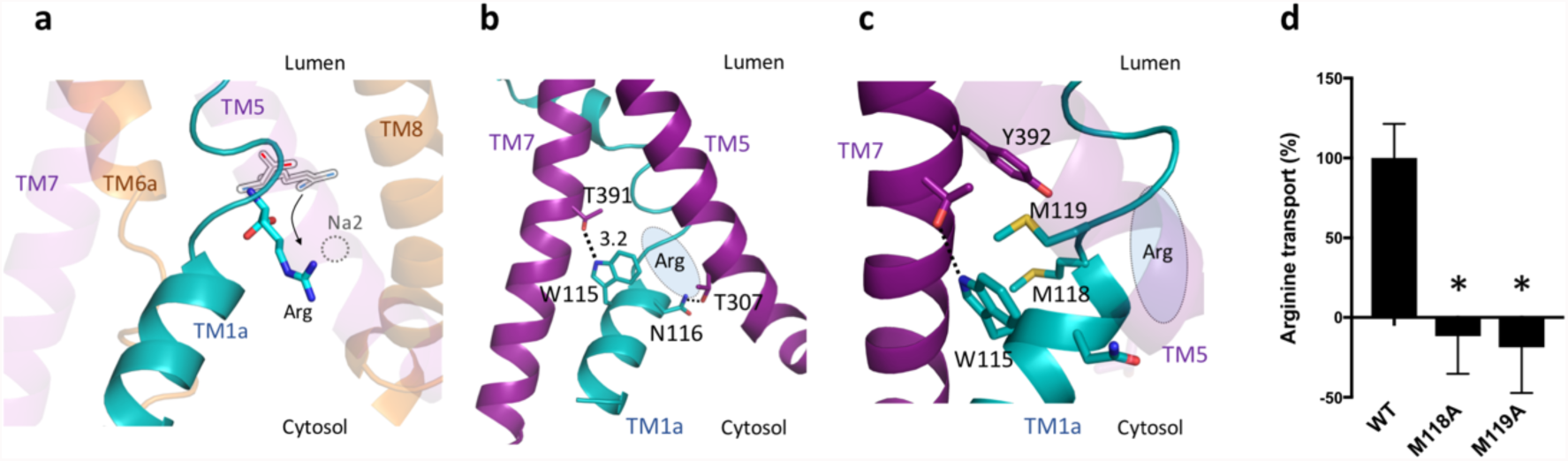
Arginine binding of SLC38A9 stabilized by TM1a which in anchored by a hydrophobic box. **a**, Arg bound in SLC38A9 (blue stick) is pointing toward the cytosol, placing the guanidinium group between TM1a and TM8. A putative Arg (grey lined hollow stick) at the central site is taken from AdiC (3L1L)^23^ after structural alignment. Superposition of SLC38A9 and vSGLT (3DH4)^25^ reveals that the sodium ion (dashed circle) at the Na2 site of the inward-facing vSGLT is located similarly to the positive guanidinium group of Arg in SLC38A9. Structural alignment is performed in PyMOL. AdiC to SLC38A9; RMSD=3.57 Å, Cealign for 72 residues. vSGLT to SLC38A9; RMSD=4.00 Å, TMalign for 168 residues. Arrow shows the putative translocation of Arg during transport. **b**, Polar interactions stabilize TM1a (cyan) between TM5 and TM7 (purple). Bonds between TM1a to TM5 and TM7 are shown as dashed lines. **c**, Hydrophobic box between TM1a and TM7. Met 118 and Met 119 are confined by Trp 115 and Tyr 392. **d**, Proteoliposome-based transport assay for arginine. M118A and M119A mutants show significant deficiency of arginine transport compared to wild-type SLC38A9. Control measurements for protein-free liposomes were subtracted from the measurements of the protein-reconstituted liposomes. Percent transport is calculated by normalizing the mean of WT to 100 %. Bar graph represents mean ± SEM of triplicate measurements. (unpaired t-test, *, p<0.05)

Like several other inward-open structures, electron densities for sodium ions are missing in the cytosol-open state of SLC38A9. Two putative sodium sites (Na1 and Na2) are identified by structural alignment with LeuT and vSGLT, coordinated by M118(O), S122, A357(O), Y392 for Na1 and N116, M119(O), Q438, T441 for Na2 (Fig. S6). Surprisingly, the sodium site Na2 in SLC38A9 was occupied by the positively charged group of bound arginine (Fig. 3a). The Na2 occupation may result in a discontinuous release of arginine in SLC38A9 transport cycle before recovering to the sodium loaded conformation. Indeed, when switching from outward-to inward-open state, the sodium ion at Na2 site is predicted to enter a metastable state and initiate the substrate release coordinated by conformational changes of TM1^25^. Thus, a similar metastable state could be expected when the sodium ion is replaced by an arginine in SLC38A9.

In the present structure, the anchoring of TM1a to TM5 and TM7 stabilizes SLC38A9 in this intermediate state. The intricate interaction network between TM1a, TM5 and TM7 was found following the bound arginine (Fig. 3b). A salt bridge is formed between Asp 116 and Thr 307. While located next to Asp 116, Trp 115 is bonded to Thr 391. Together, Trp 115 and Asp 116 marks the beginning of this anchor network on TM1a in the cytosol-open structure (Fig. 3b). The TM1a anchor suspends the helical segment of TM1a from its cytosolic end and contains a hydrophobic box formed by Met 118 and Met 119 sandwiched between W115 and Tyr 392 (Fig. 3c). This hydrophobic box is immediately followed by the unwound region on TM1, suggesting that TM1a anchor may cause Met 119 to expose its carbonyl oxygen to disrupt the alpha helix. With the conserved WNTMM motif, a restrained TM1a in this conformation is likely to be important during transport by SLC38A9 homologs. Consistent with the structural insights mentioned above, a mutation N128A in the human homolog (hSLC38A9) has been shown to decrease transport activity^3^. While sequence alignment shows that human Asp 128 corresponds to Asp 116 in the zebrafish homolog (Fig. S1), disturbing Asp 116 in the WNTMM motif on TM1a is believed to impair the anchoring network along with sodium coordination, which can consequently undermine transport.

Mutations M118A and M119A in WNTMM motif abolished the transport of arginine, suggesting that the large non-polar side chains at this position are functionally necessary (Fig. 3d). Presumably, the two “methionine fingers” insert into the hydrophobic box during conformational changes and draw TM1a to open at the cytosol. Tracing up from TM1a, TM1 and TM6 draw close at the GTS motif (Fig. S7) toward the luminal end. In the cytosol-open state of SLC38A9, TM1a alone mediates arginine binding and forms a stabilized conformation. To achieve the same task, other LeuT-like transporters (for example vSGLT and CaiT) would use both TM1 and TM6 to retain its substrates in the inward-open state, involving the GTS motif at disrupted regions (Fig. S7)^21,26^.

The conformational changes of TM1 are critical for SLC38A9 using the central binding site (at Thr 121) for substrate uptake and efflux^2^. Moreover, because of the pivotal role of SLC38A9 as a transceptor in modulation of mTORC1^10^, each conformational state may have a designated purpose to carefully regulate the downstream signaling of mTORC1^2,27–29^. The structure of SLC38A9 with an arginine at the TM1a site allows us to propose a schematic model of arginine release by SLC38A9 in three states (Fig. 4). It is known that arginine is transported with sodium ions in SLC38A9. In the active transport state (putative State1), arginine would bind at the central site and would be coordinating a sodium ion at the Na1 site. The α-carboxylate group of arginine is supposedly bound by TM1 and TM6 at the hinge region. The guanidinium group of arginine is stabilized by TM3. When open to cytosol (State2), the sodium ion bound in Na2 site is destabilized and released after the conformational change of TM1a. TM1a is arrested by two polar interactions, W115-T391 and N116-T307, with TM5 and TM7. The TM1a anchor renders a new binding site next to TM1a for arginine, connecting the central binding site to the cytosol. The guanidinium group of arginine comes in proximity of Na2 site and occupies this location after the sodium ion was released between TM1a and TM8. Arginine would need to complete a rotation of ~90° about the α-amino group to reach the TM1a binding site from the central site. When arginine is released (State 3), TM1a binding site is emptied by solvent diffused in the cytosolic vestibule. TM1a becomes unstable, and the TM1a anchor is broken from TM5 and TM7.

**Figure 4.**
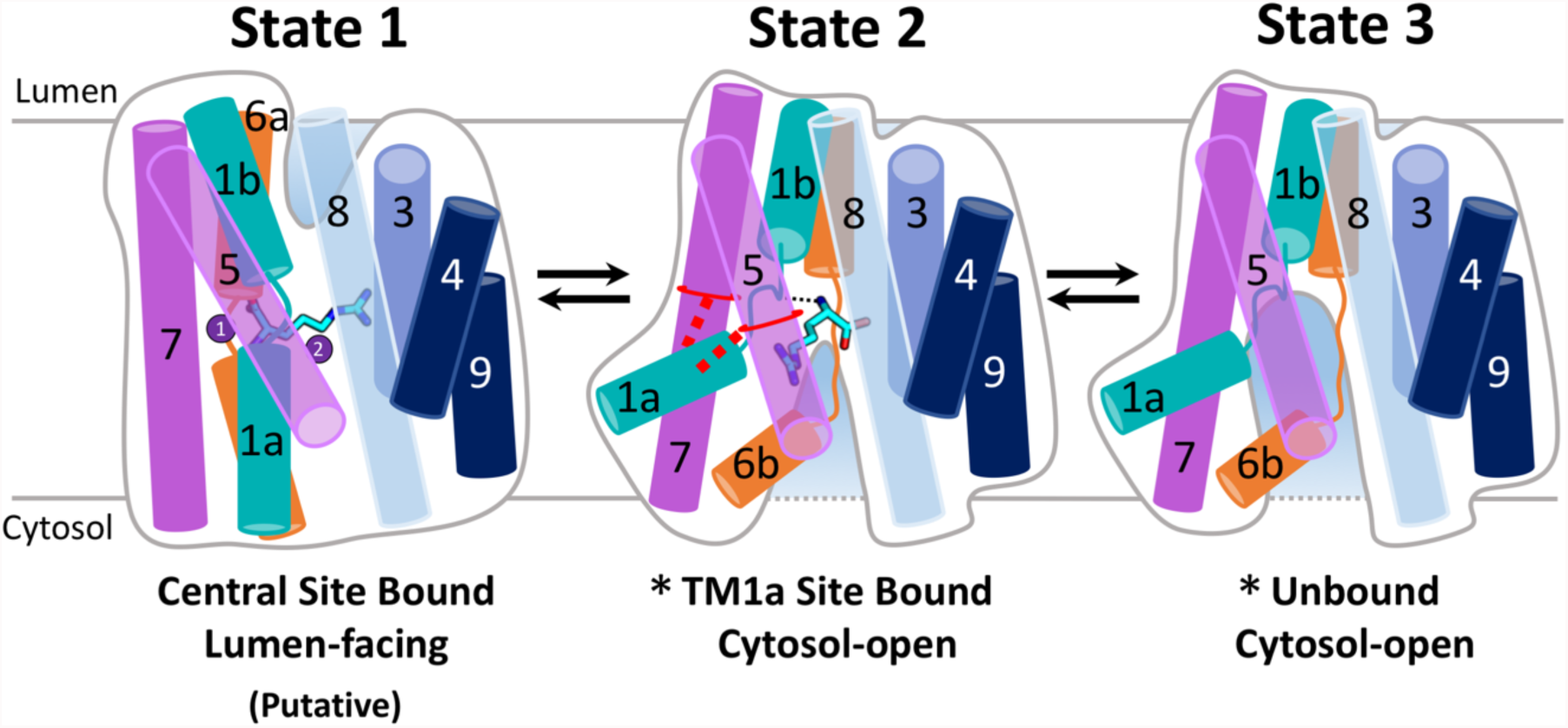
Proposed cytosolic release of Arg by SLC38A9. State 1, A putative lumen-facing conformation, Arg is bound at central site and coordinated by Na ion at Na1 site. α-carboxylate group of Arg is bound to TM1 and TM6 at the disrupted region. Guanidinium group of Arg is pointing to TM3. The two predicted Na1 and Na2 sites (purple sphere 1 and 2) are located by aligning to LeuT (3TT1) and vSGLT (3DH4) (Fig. S3)^19,25^. State 2, Arg is bound at the TM1a binding site as elucidated in the crystal structure. TM1a is anchored by a pair of residues on TM5 and TM7, rendering a negatively charged binding site for Arg and connecting the central site to cytosol. Guanidinium group of Arg occupies Na2 site and displaces sodium ion, pointing to TM5. State 3, Arg is released from TM1a binding site. SLC38A9 is emptied by buffer diffused into the cytosolic vestibule. TM1a anchor is weakened and cannot maintain interactions with TM5 and TM7. *Two copies of SLC38A9 are found in the asymmetric unit of the crystals, revealing Arg-bound and Arg-free cytosol-open states within the same crystal form.

Here we described the first structure of SLC38A9 and discovered a novel TM1a-arginine-binding site in the transporter captured in the cytosol-open state. The TM1a binding site consists of an anchor with two critical methionine fingers inserted into a hydrophobic box. Movement of the TM1a anchor is proposed to lead to an intermediate state during arginine uptake or release, which may regulate amino acid transport and modify the transport efficiency of the transporter in the presence of arginine. Importantly, the intermediate state of SLC38A9 described in this study suggests that arginine binding could enable the N-terminus of SLC38A9 to bind with Rag GTPases and Ragulator by moving the TM1a which directly links the transmembrane domain and the N-terminal domain of SLC38A9. Once Rag GTPases and Ragulator bind to SLC38A9, the downstream mTORC1 signaling could then become activated^3,5^. However, a complete structure of SLC38A9-Ragulator-Rag GTPases complex captured at different conformational states will be needed to elucidate the precise process by which this transporter activates this pathway and to advance our understanding toward the lysosomal amino acid transport and its modulation of mTORC1 signaling pathway. The understanding of amino acid sensing and transport by SLC38A9 will further assist in drug development against the deregulated mTORC1 activation and lysosomal homeostasis related disorders^30,31^.

## METHODS SUMMARY

SLC38A9 from *Danio rerio* and its mutants were overexpressed in *Spodoptera frugiperda* Sf-9 cells. Fab antibody fragments against the luminal epitope of SLC38A9 were generated and purified as described in Methods. The SLC38A9-Fab complex was purified in the presence of 0.2% (w/v) n-decyl-β-D-maltoside (Fig. S8) and crystallized in the following condition, 30% PEG 400, 0.1 M N-(2-Acetamido) iminodiacetic acid (ADA) pH 7.2~7.4, 0.35 M Lithium Sulfate. Crystals were grown or soaked in the same crystallizing condition supplemented with 5-20 mM L-arginine. Diffraction data sets of all crystals were collected at the Advanced Photon Source (NE-CAT 24-ID-C and ID-E). Data processing and structure determination were performed using RAPD, PHENIX and Coot. Detailed methods can be found in the Methods accompanying this manuscript.

## Acknowledgements

We thank D. Cawley for development and production of monoclonal antibodies. We thank K. Rajashankar and the staff in NECAT for their support with X-ray data collection. We thank D. Casio and J. Hattne for discussions over X-ray data collection and structural determination. We thank L. Shao and S. Liu for critical reading of the manuscript. This work is based upon research conducted at the Northeastern Collaborative Access Team beamlines, which are funded by the National Institute of General Medical Sciences from the National Institutes of Health (P41 GM103403). The Pilatus 6M detector on 24-ID-C beam line is funded by a NIH-ORIP HEI grant (S10 RR029205). This research used resources of the Advanced Photon Source, a U.S. Department of Energy (DOE) Office of Science User Facility operated for the DOE Office of Science by Argonne National Laboratory under Contract No. DE-AC02-06CH11357. Research in the Gonen laboratory is funded by the Howard Hughes Medical Institute.

## Author Contributions

H.-T.L., J.M. and T.G. designed the project. H.-T.L. and J.M. performed experiments including protein preparation, antibody screening, crystallization and data collection. H.-T.L. performed model building and refinement. S.S.M. and H.-T.L. performed the radioligand uptake assay. H.-T.L., J.M. and T.G. participated in data analysis and figure preparation. H.-T.L. and T.G. wrote the manuscript.

## Author Information

Coordinates and structure factors were deposited in the Protein Data Bank (PDB accession code 6C08). Reprints and permissions information is available at www.nature.com/reprints. The authors declare no competing financial interests. Readers are welcome to comment on the online version of the paper. Correspondence and requests for materials should be addressed to T.G.

---

**Table S1.** Data Collection and refinement statistics table.

|  | SLC38A9-11D3(Native) * | SLC38A9-11D3(Se-Met) |
| --- | --- | --- |
| **Data collection** |  |  |
| Space group | P 1 21 1 | P 1 21 1 |
| Cell dimensions |  |  |
| a, b, c (Å) | 137.27, 82.83, 159.37 | 136.163, 82.795, 158.922 |
| $\alpha$, $\beta$, $\gamma$ (°) | 90, 100.13, 90 | 90, 100.005, 90 |
| Wavelength | 0.9791 | 0.9791 |
| Resolution (Å) | 3.1 | 3.358 |
| R<sub>meas</sub> | 0.127 (1.642) | 0.07465 (1.812) |
| Mean I/sigma(I) | 8.3(1.1) | 12.71 (0.86) |
| Completeness (%) | 99.7 (98.1) | 96.98 (81.17) |
| Redundancy | 8.3 (8.5) | 3.4 (3.2) |
| CC1/2 | 0.997 (0.6) | 0.999 (0.347) |
| **Refinement** |  |  |
| Resolution (Å) | 156.5 - 3.17 (3.283 - 3.17) |  |
| No. reflections | 58098 (5909) |  |
| CC1/2 | 0.997 (0.492) |  |
| R<sub>work</sub>/R<sub>free</sub> | 0.2680 (0.3580) /0.2845 (0.3721) |  |
| No. atoms |  |  |
| Protein | 12332 |  |
| Ligand/ion | 1 Arg |  |
| Average B-factor (Å<sup>2</sup>) | 114.00 |  |
| SLC38A9 | 141.23 |  |
| 11D3-Fab | 88.75 |  |
| Arg | 147.08 |  |
| R.m.s. deviations |  |  |
| Bond length (Å) | 0.006 |  |
| Bond angle (°) | 1.36 |  |
| Ramachandran |  |  |
| Outliers (%) | 4.36 |  |

\*merged from two crystals

**Figure S1.**
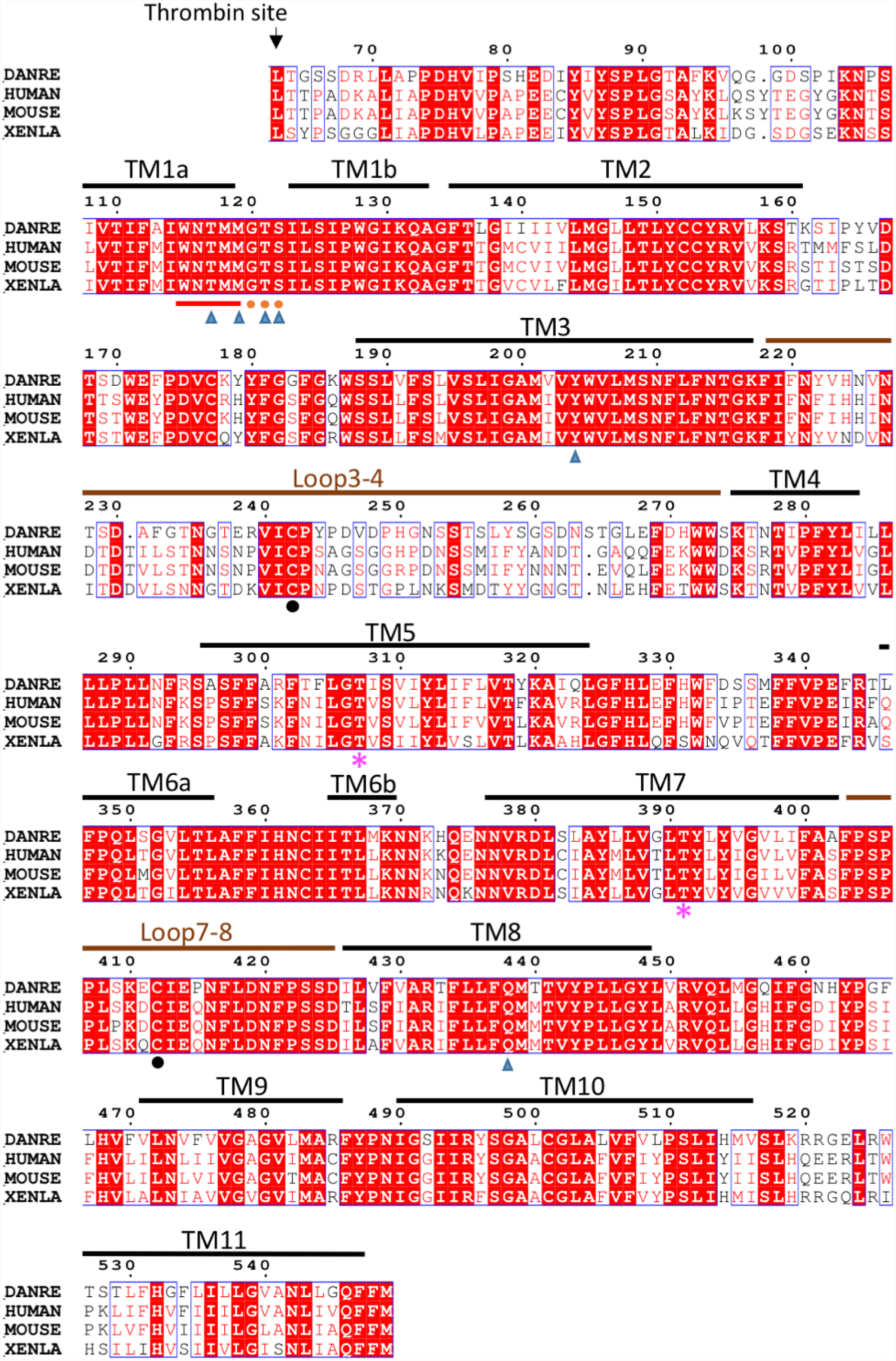
Sequence alignment of SLC38A9 and its homologs. Full length protein sequence alignment of Primary accession number in order: Q08BA4 (zebrafish, DANRE), Q8NBW4 (human), Q8BGD6 (mouse), Q6DFK0 (frog, XENLA). Black lines, transmembrane regions; brown lines, loops bound to Fab; red underline, WNTMM anchor motif; orange circles, GTS conserved motif; magenta asterisks, residues bonded with TM1a; blue triangles, residues involved in Arg binding; black circles, disulfide bond between Loop3-4 and Loop7-8.

**Figure S2.**
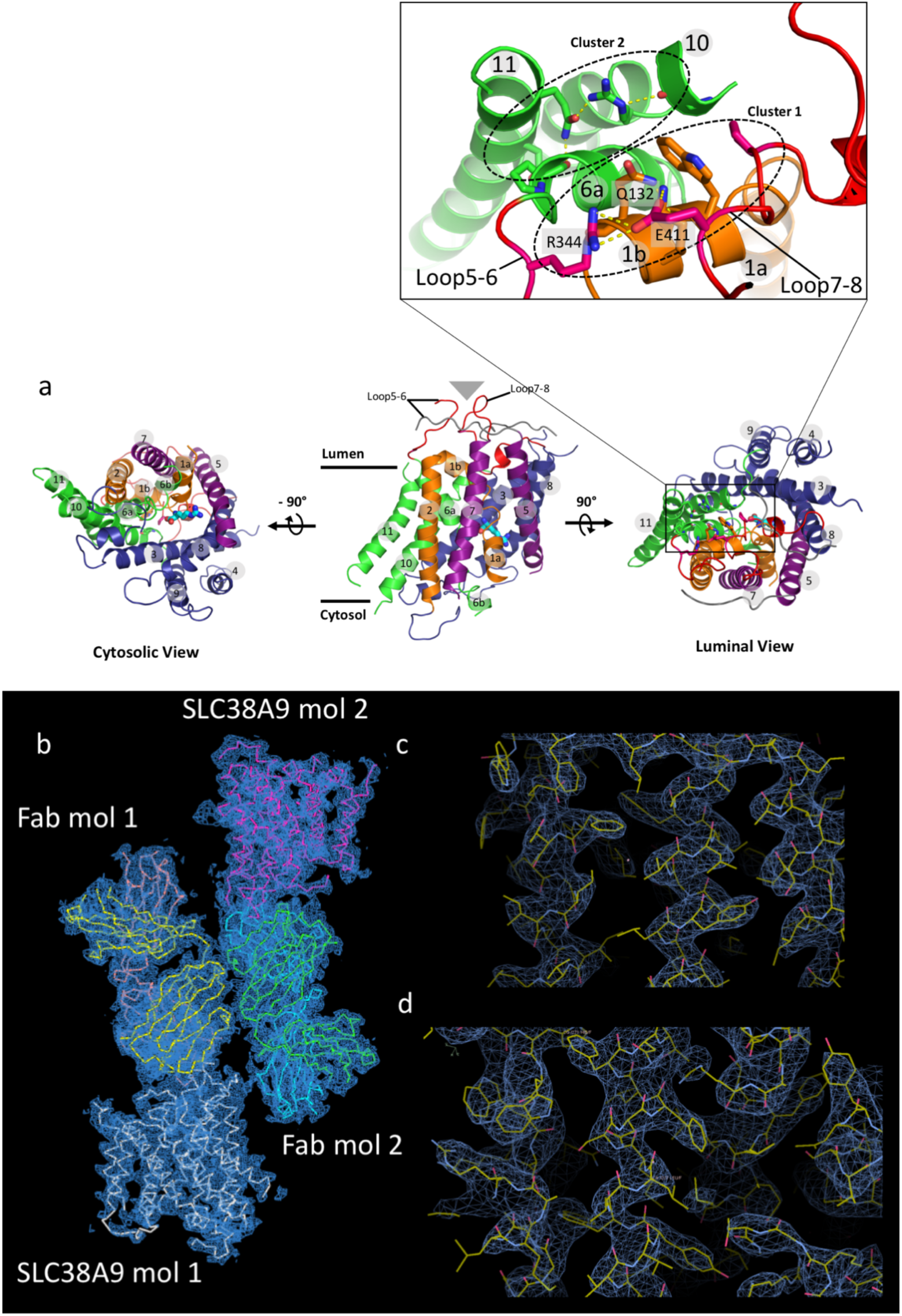
Overall structure of SLC38A9. **a**, Central panel, side view in plane of lysosomal membrane. Position of Fab fragment bound on the luminal side is shown by grey triangle above the luminal loops. Left panel, cytosolic view showing the vestibule at cytosolic side. Right panel, luminal view. At luminal side, residues on TM1b, Loop5-6, Loop7-8 are grouped in cluster 1(W128, K131, Q132, R344, E411, P415) and TM6a, TM10, TM11 are grouped in cluster 2 (P348, G491, R495, N542, Q546). Enlarged window from the luminal view encompasses luminal gating cluster 1 and cluster 2. **b**, The electron density of the determined asymmetric unit of SLC38A9-Fab crystals. Each asymmetric unit contained two SLC38A9 molecules (labeled SLC38A9 mol 1 and mol 2) as well as two Fab fragments (labeled Fab mol 1 and Fab mol 2). **c-d**, Examples of the quality of the electron density maps in the determined structure with the fitting of the model of SLC38A9-Fab.

**Figure S3.**
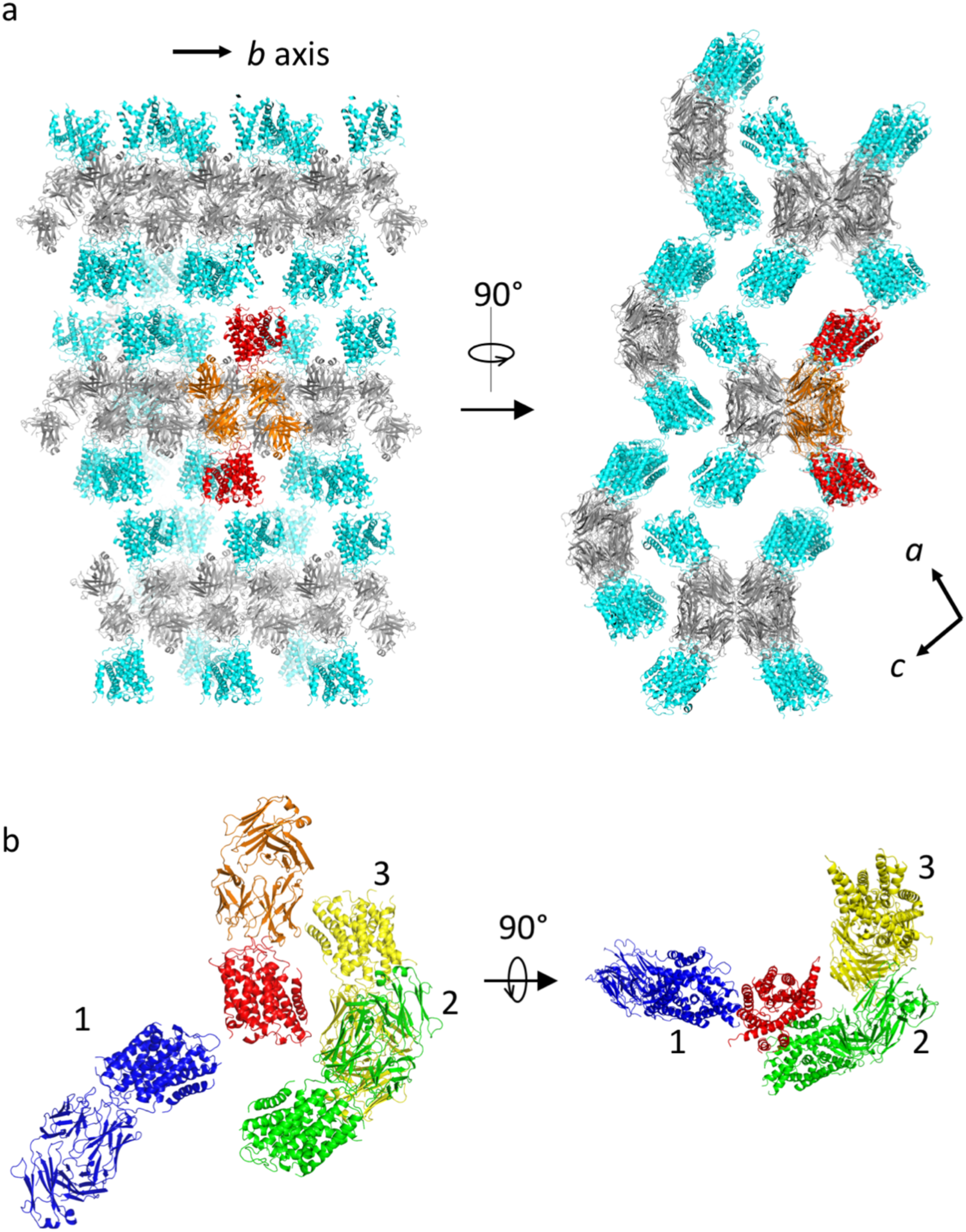
Crystal packing and asymmetric unit of the SLC38A9-Fab complex. **a**, Crystal packing showing SLC38A9-Fab complex lattice. Fab (grey) stacks tightly along the crystallographic *b* axis, and are connected by SLC38A9 (cyan) layers in the crystallographic *ac* plane in a propeller-like head-to-side manner. One asymmetric unit is selected to show the building block that is composed of two Fab (orange) and two SLC38A9 (red) molecules. **b**, Interactions between SLC38A9 and adjacent Fab fragments. One SLC38A9 (red) makes contacts with four other molecules. The biologically functional contact is between the luminal loops of SLC38A9 (red) and the complementary determining regions (CDRs) of the Fab (orange). The three other contacts, which appear to be crystal contacts and non-specific, occur between Loop2-3 (red) and TM5 (blue 1), and between TM3, TM10 (red) and a groove shaped from the two adjacent Fab fragments (green 2 and yellow 3).

**Figure S4.**
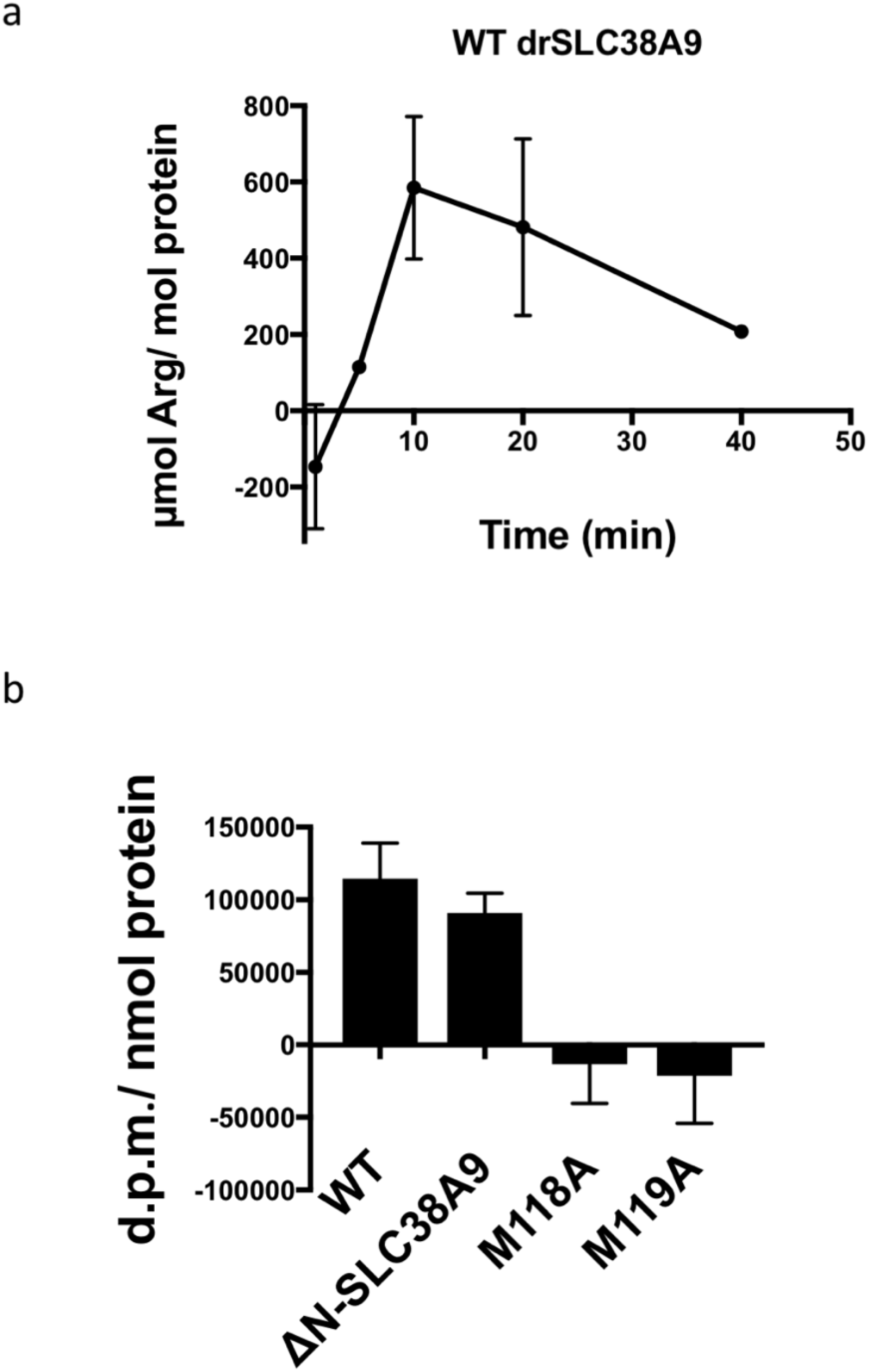
Proteoliposome-based transport assay **a**, Time course of [^3^H]-Arg uptake by wild-type drSLC38A9. [^3^H]-Arg uptake was followed for 40 minutes. Amount of retained [^3^H]-Arg saturated after 10 minutes of incubation with 0.5µM of external [^3^H]-Arg. Judging from this result, we selected a time point of 5-minute incubation to compare the transport ability of different protein constructs. Error bar, standard error of the mean (s.e.m.) of triplicate measurements. **b**, d.p.m./ nmol protein measurements at 5 minutes of incubation for wild-type, ΔN-SLC38A9, M118A and M119A mutants. Error bar, standard error of the mean (s.e.m.) of triplicate measurements. One-sample t-test was used to compare M118A and M119A to value 0, which confirms that the two mutants are deficient in arginine transport *in vivo* at p values of 0.7072 and 0.6315 for M118A and M119A respectively.

**Figure S5.**
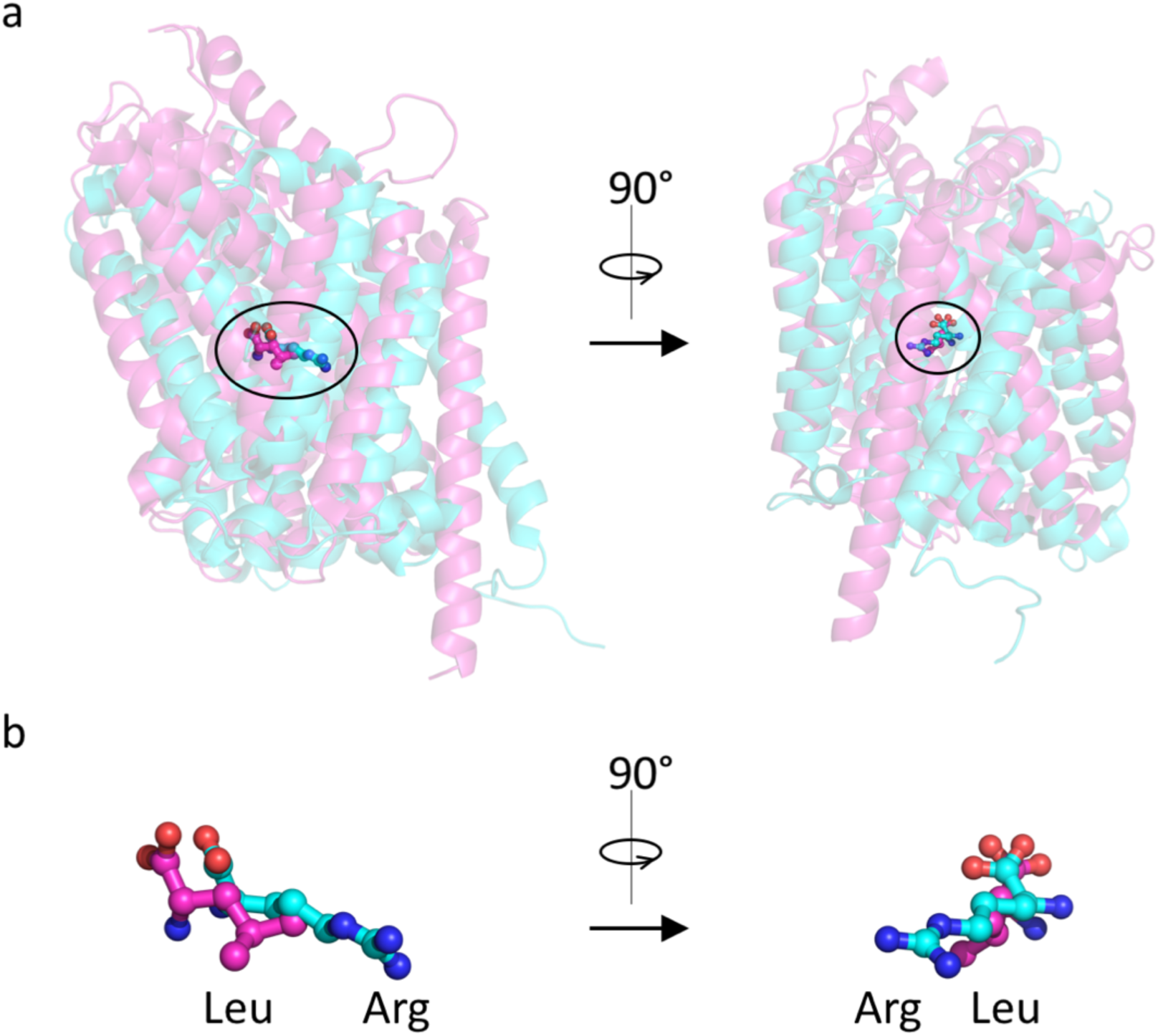
The structural alignment of substrate-bound AdiC and LeuT. **a**, AdiC (3L1L)^23^ in cyan and LeuT (2A65)^32^ in magenta is aligned in PyMOL, with the whole chain calculated by Cealign for 360 residues (RMSD =4.94), revealing the similar central site locations (solid circle). b, Close up view of the bound Arg from AdiC (cyan) and Leu from LeuT (magenta). The common groups of Arg and Leu are overlayed in between TM1 and TM6, while both side groups of Arg and Leu are pointing to TM3.

**Figure S6.**
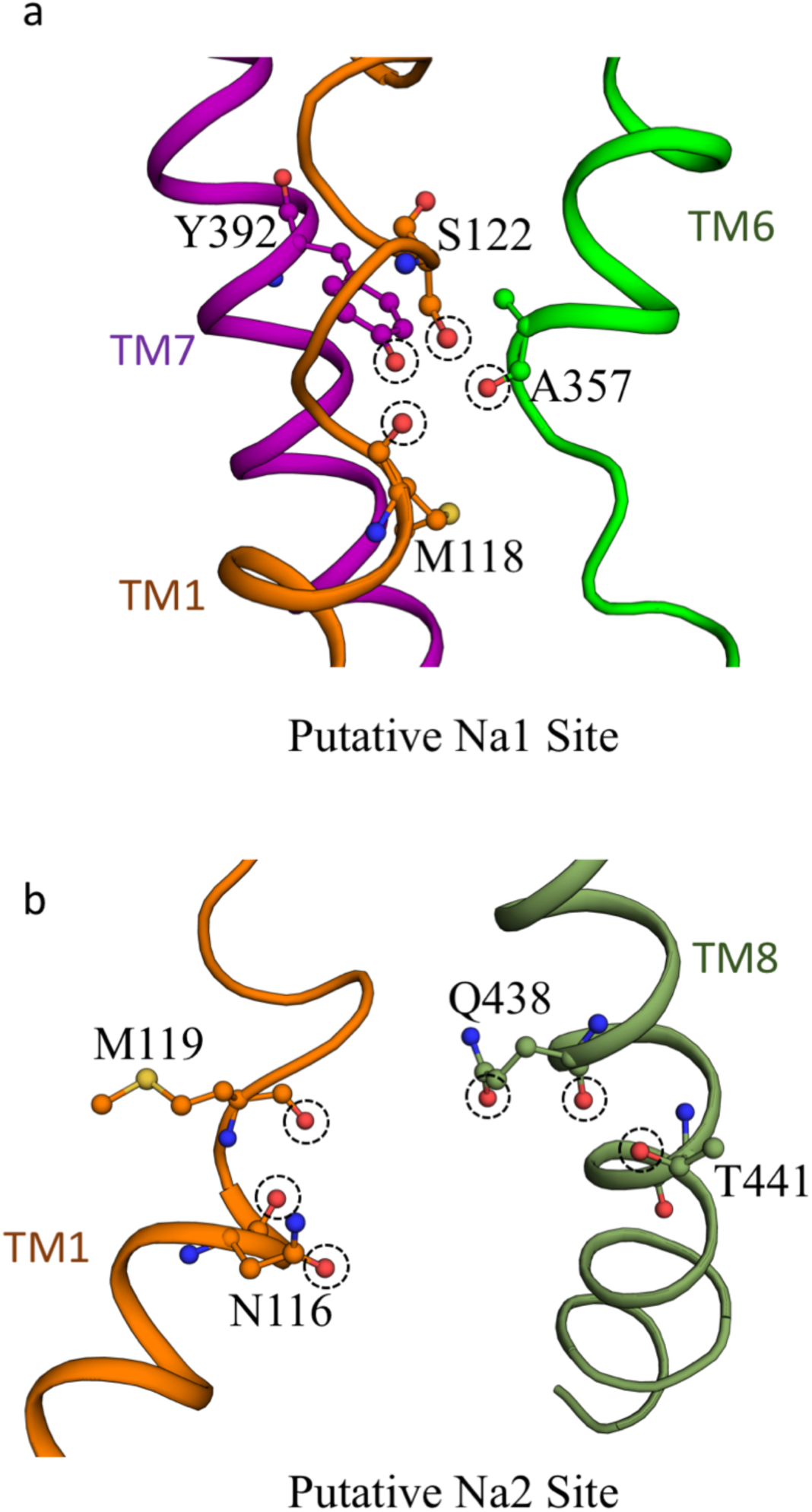
Putative sodium binding sites. **a**. Predicted sodium ion binding site Na1 by structural alignment to LeuT (PDB 3TT1, RMSD=3.92, TMalign for 185 residues). Sodium ion at Na1 site is coordinated by four oxygens (circled by dash lines) from M118(O), S122, A357(O), Y392. **b.** Predicted sodium ion binding site Na2 by structural alignment to vSGLT (PDB 3DH4, RMSD=4.00, TMalign for 168 residues). In the current structure, sodium ion at Na2 site is displaced by arginine. However, four residues, N116, M119, Q438, T441 are likely to become electron donors when bound to sodium ion (circled by dash lines).

**Figure S7.**
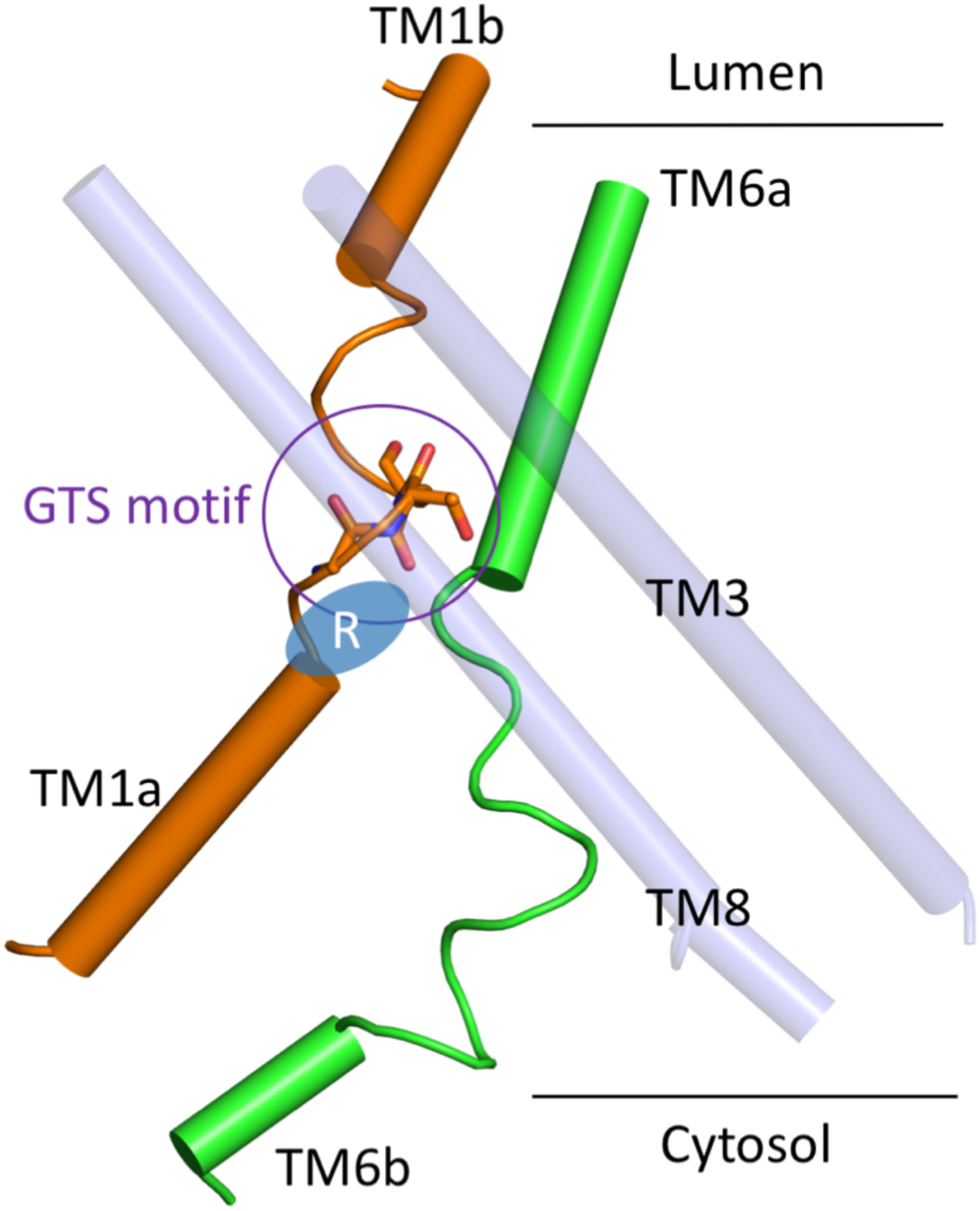
GTS motif on TM1 and Arg in TM1a binding site. The gap space between TM1 and TM6 closes up after GTS motif. Position of Arg is shown in blue oval. Recognition of Arg at this location is independent of TM6.

**Figure S8.**
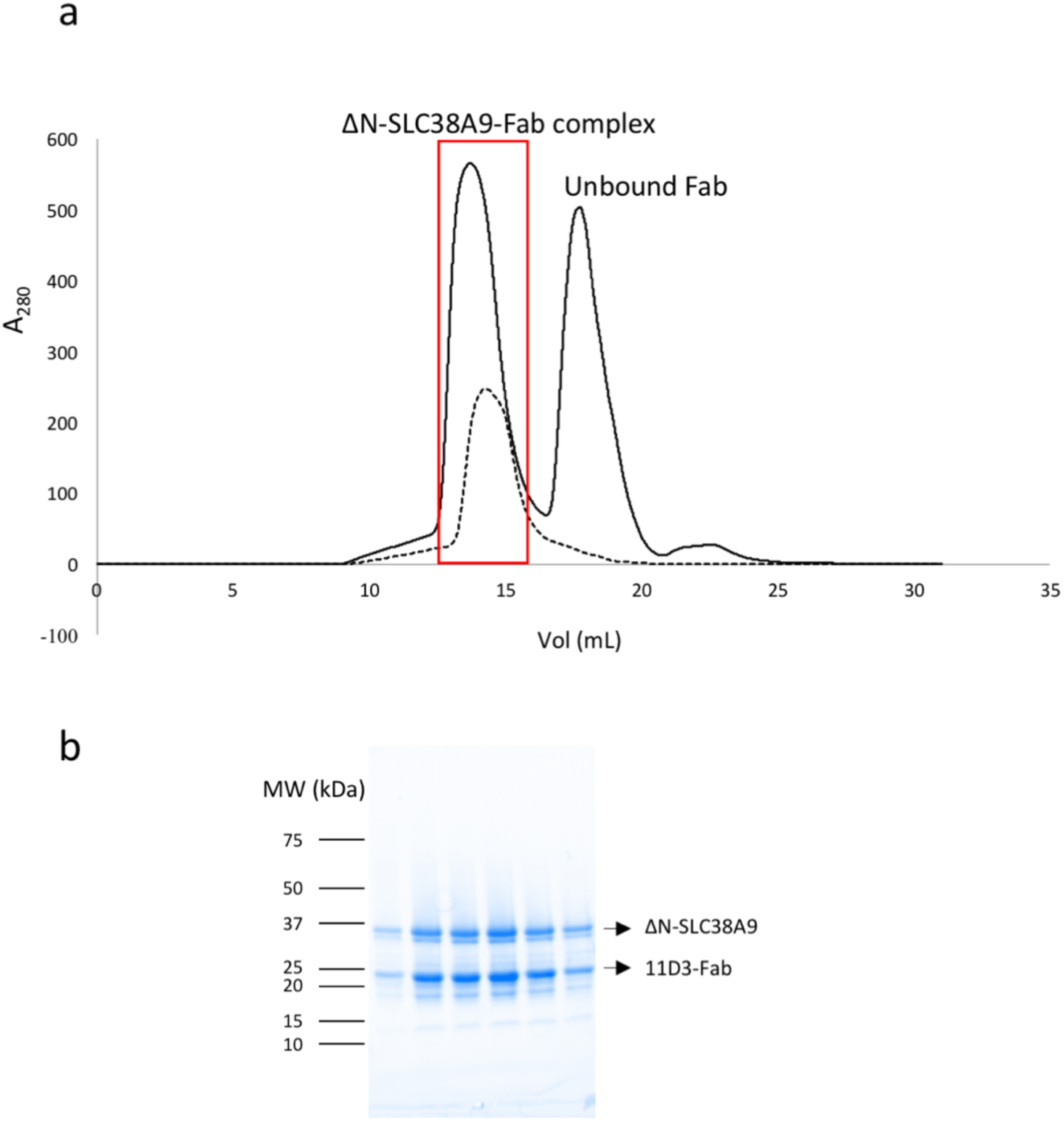
Elution profile of the ΔN-SLC38A9-Fab assembly. **a**, Solid line represents the elution trace of ΔN-SLC38A9-Fab complex and unbound Fab. Dash line is the elution trace of pure ΔN-SLC38A9. Apparent peak shift was observed for the formation of ΔN-SLC38A9-Fab complex. **b**, Fractions selected in the red box shown in **a** was sampled and analyzed on SDS-PAGE before pooled and concentrated for crystallization.

## METHODS

### SLC38A9 cloning, expression and purification

Gene encoding N-terminal truncated SLC38A9 (accession number Q08BA4), ΔN-SLC38A9, was cloned into pFastBac-1 vector (Invitrogen) with a N-terminal 8 X His-tag and a thrombin cleavage site. Four mutations (N227Q, N235Q, N252Q, N263Q) were introduced at the glycosylation sites. Plasmids were transformed in DH10bac for preparation of bacmids. Recombinant baculovirus was generated and used in transfection following the protocol provided in Bac-to-bac Baculovirus Expression System. ΔN-SLC38A9 was overexpressed in *Spodoptera frugiperda* Sf-9 insect cells, which was harvested 60h post-infection. Cell pellets was resuspended in lysis buffer containing 20 mM Tris (8.0), 150 mM NaCl supplemented with protease inhitior cocktails (Roche). 40 homogenizing cycles were then carried out to break cells on ice, followed by centrifugation at 130,000 × g for 1h. Pelleted membrane was resuspended and washed in high salt buffer containing 1.6 M NaCl and 20 mM Tris (8.0) and centrifuged again for 1h at 130,000 × g. The pelleted membrane was frozen in liquid N_2_ and stored in −80°C until further use. To purify ΔN-SLC38A9, membrane pellet was solubilized in 2 % n-dodecyl-b-D-maltopyranoside (DDM), 20 mM Tris (8.0), 150 mM NaCl, 5 % glycerol, 0.2 % Cholesteryl Hemisuccinate Tris Salt (CHS) for 4 hours at 4°C, followed by 1h centrifugation at 130,000 × g. 20 mM Imidazole (8.2) was added to the supernatant before incubation with TALON beads for 16 h at 4°C. ΔN-SLC38A9 bound beads were washed by 6 column volumes of 20mM Imidazole, 20 mM Tris (8.0), 500 mM NaCl and 0.1 % DDM. The resins were then equilibrated in buffer 20 mM Tris (8.0), 150 mM NaCl, 0.4 % decyl-b-D-maltoside (DM) and 0.02 % DDM. At 4°C, 8 × HisTag was removed by in-column thrombin digestion overnight at enzyme:protein molar ratio of 1:500. The cleaved ΔN-SLC38A9 was collected in flow-through and was flash frozen in liquid N2 and stored in −80°C until use. SeMet substituted ΔN-SLC38A9 was overexpressed in Sf-9 cells using the same procedures described above for native protein except that 100 mg L^−1^ seleno-methionine (Acros Organics) was added to cultures during the course of 60 h post-infection. Purifying procedures of Se-Met substituted ΔN-SLC38A9 were the same as the native protein.

### Fab production and purification

Mouse IgG mono-clonal antibodies against ΔN-SLC38A9 was produced by Monoclonal Antibody Core, Dr. Daniel Cawley. 330 µg purified ΔN-SLC38A9 in the buffer containing 20 mM Tris (8.0), 150 mM NaCl, 0.02 %DDM, 0.002 % CHS was used to immunize mice in three injections. 15 × 96 well plate fusions yielded 169 IgG positive wells at a 1/30 dilution. Native and denatured ΔN-SLC38A9 were then used in ELISA to search for candidates that bind the conformational epitopes^33^, where Ni-NTA plates were used for ΔN-SLC38A9 immobilization. 35 of the 169 fusions showed significant preferences of binding against well-folded ΔN-SLC38A9. Western blot was performed to assess the binding affinity and specificity of the antibodies generated from hybridoma cell lines. Monoclonal antibody 11D3 was then purified from the hybridoma supernatants by 4-mercapto-ethyl-pyridine (MEP) chromatography. Fab was produced by papain digestion and purified in the flow-through buffer containing 20 mM NaPi (8.0), 150 mM NaCl by protein A affinity chromatography.

### Assembly of ΔN-SLC38A9–Fab complex

Purified Fab fragment of 11D3 was added to ΔN-SLC38A9 at 2:1 molar ratio, and was incubated for 4 h to form stable complexes. ΔN-SLC38A9-11D3 was concentrated by centrifugal filter vivaspin 20 at 50 kDa MWCO. The concentrated protein is further purified and underwent a detergent exchange by gel filtration, Superdex-200 size exclusion column (SEC), in buffer containing 20 mM Tris (8.0), 150 mM NaCl, 0.2 % DM. Judged by SDS-PAGE and size exclusion chromatography elution profile (Fig. S9), fractions containing appropriate ΔN-SLC38A9-11D3 complexes were pooled and concentrated to 5 mg mL^−1^ for crystallization.

### Crystallization

Initial hanging-drop crystallization assay with purified ΔN-SLC38A9 produced crystals grown in the condition of 30 % PEG400, 100 mM Tris (8.0) and 400 mM LiCl at 4°C. However, these crystals gave anisotropic diffraction to around 6 Å. Crystals showing adequate diffraction power were obtained only when ΔN-SLC38A9 was co-crystallized as a complex with Fab prepared from hybridoma cell line 11D3 (IgG2a, kappa). The best crystal, which diffracted to 3.1 Å, was obtained from the condition of 26-30 % PEG400, 100 mM ADA (7.2) and 350 mM Li_2_SO_4_ at 4°C. Before data collection, the crystals were soaked in the same crystallizing solution containing 30 % PEG400 and 20 mM arginine pH 7.2 for 1 h, and were rapidly frozen in liquid N_2_. Se-Met crystals were grown and harvested in the same manner as the native crystals.

### Data collection and structure refinement

X-ray diffraction datasets were collected at the Advanced Photon Source (Argonne National Laboratory, beamline 24-IDC and 24-IDE), and processed in the online server RAPD, which uses XDS and CCP4 suite package for integrating and scaling to resolutions of 3.1 Å (Native) and 3.4 Å (Se-Met). Antigen-binding fragments (Fab) from PDB 1F8T^34^ was used in the initial molecular replacement as the search model. The Se-Met dataset was then phased by single anomalous dispersion in Phenix^35^ using differences from 23 Se atoms at lambda = 0.9791 Å and the two Fab fragments previously placed using Phaser as a partial model (MRSAD)^34^. Helices were manually placed in the density-modified map and extended using Coot^36^. Subsequent cycles of density modifications, model building and refinement were carried out in Phenix and Coot until structure completion. The final model contains two molecules of ΔN-SLC38A9 (residues 108–549) and two pairs of the heavy-light chain of Fab in an asymmetric unit. Data collection and refinement statistics are presented in Supplementary Table S1.

### Preparation of full-length SLC38A9 and ΔN-SLC38A9 proteoliposomes

Full-length and N-terminal truncated SLC38A9 were expressed and purified as described above. Liposomes were prepared by resuspending thin films of 35mg/mL egg phosphatidylcholine (egg-PC) in buffer A (20 mM MES pH7.0, 100 mM NaCl and 1 mM DTT), followed by extrusion through membranes with pore size of 0.1 µm. Triton X-100 was added to the extruded liposomes at 10:1 (w:w) lipid:detergent ratio. Volume was then adjusted to final concentration of 14 mg egg-PC/ mL with buffer A and incubated for 1h, followed by reconstituting SLC38A9 or ΔN-SLC38A9 at protein-to-lipid ratio of 1:400 (w:w) for 2h. Detergents were removed by SM2 Bio-Beads (Bio-Rad) added to the protein-lipid mix and rotated overnight at 4°C. Next day, proteoliposomes were collected, aliquoted and stored at –80 °C.

### Radioligand uptake assays

Proteoliposomes of reconstituted SLC38A9, ΔN-SLC38A9, M118A and M119A mutants were thawed on ice. Transport reactions were initiated by adding 0.5 µM L-[^3^H]-arginine (American Radiolabeled Chemicals, Inc) to 100 µL aliquots of proteoliposomes. A negative control of protein-free liposomes was carried out in parallel to experiment groups. At various time points, proteoliposomes were filtered, washed by 1mL of buffer A, and collected on 0.22µm GSTF nitrocellulose membranes. 10 mL of scintillation fluid was then added to each filter in a vial and counted. A time course profile indicates that the retained radio-ligands reached saturation in 10 min (Fig. S4). Measurements at 5 min of the arginine uptake were used to establish the transport comparisons between various constructs of SLC38A9, normalized to that of full-length wildtype SLC38A9. Nonspecific adsorptions of L-[^3^H]-arginine by egg-PC liposomes were subtracted from the experiment measurements. Each of the experiments were repeated three times. Raw data of counted radioactivity is shown in Fig. S4. Statistical comparisons were made from unpaired t-test calculations, in which the P value cutoff is set to 0.05 for significance level in GraphPad Prism Version 7.0c.

All figures in this paper were prepared with PyMOL v1.8.6.0 ^37^and assembled in Microsoft PowerPoint v15.18. Figure. S1 was prepared using the program Clustal Omega^38^ for alignments and ESPript 3.0 ^39^ for styling.

